# GEFs and Rac GTPases control directional specificity of neurite extension along the anterior-posterior axis

**DOI:** 10.1101/052019

**Authors:** Chaogu Zheng, Margarete Diaz-Cuadros, Martin Chalfie

## Abstract

Although previous studies have identified many extracellular guidance molecules and intracellular signaling proteins that regulate axonal outgrowth and extension, most were conducted in the context of unidirectional neurite growth, in which the guidance cues either attract or repel growth cones. Very few studies addressed how intracellular signaling molecules differentially specify bidirectional outgrowth. Here, using the bipolar PLM neurons in *C. elegans*, we show that the guanine nucleotide exchange factors (GEFs) UNC-73/Trio and TIAM-1 promote anterior and posterior neurite extension, respectively. The Rac subfamily GTPases act downstream of the GEFs; CED-10/Rac1 is activated by TIAM-1, whereas CED-10 and MIG-2/RhoG act redundantly downstream of UNC-73. Moreover, these two pathways antagonize each other and, thus, regulate the directional bias of neuritogenesis. Our study suggests that directional specificity of neurite extension is conferred through the intracellular activation of distinct GEFs and Rac GTPases.

**Significance Statement:** Most previous studies on intracellular signaling during neurite guidance were performed in the context of unidirectional neurite growth. They could not address the molecular basis of directional outgrowth of multiple neurites mainly because of the lack of a good model system. Using a pair of bipolar neurons in the nematode *Caenorhabditis elegans*, we found that distinct sets of intracellular molecules are required for neurite extension towards the anterior and the posterior. Moreover, signaling pathways that promote neurite extension in different directions antagonize each other to achieve balanced growth. Therefore, our study offers an *in vivo* example for a long-standing concept that spatially selective activation of intracellular signaling molecules could enable a diverse range of neuronal growth patterns.

## Introduction

During development, axons and dendrites emerge from neuronal cell bodies and navigate towards their target. Previous studies of neurite guidance have focused on identifying both extracellular guidance molecules and their receptors (reviewed in 1) and intracellular effectors that regulate cytoskeleton dynamics and growth cone movement (reviewed in 2). Although several signaling cascades connect the activation of various ligand-bound receptors to the remodeling of the actin and microtubule cytoskeletons, they do so through similar second messengers (i.e. kinases, phosphatases, GTPases; 3, 4) and, thus, do not address how the directed outgrowth of multiple neurites occurs. Presumably distinct intracellular pathways mediate neurite outgrowth and extension in different directions, but what these pathways are remains unclear.

One potential means of regulating differential outgrowth is the use of the many members of Rho family of small GTPases [Rac (including Rac1, Rac2, Rac3, and RhoG), Cdc42, and Rho] and the guanine nucleotide exchange factors (GEFs) that activate these GTPases (5). Studies using *in vitro* cultured mammalian neurons suggest that Rac1 (activated by the GEFs Tiam1 and Dock180), RhoG (activated by the first GEF domain of Trio), and Cdc42 promote axonal outgrowth and extension by regulating the actin and microtubule cytoskeleton (6, 7). *In vivo* studies in Drosophila and *C. elegans* suggest that Rac GTPases have overlapping functions in the control of axon growth and guidance and that the Trio GEF is essential for Rac activities in the nervous system (8–10). In contrast, Rho and its downstream effector ROCK negatively regulate axon growth in mammals (11, 12) and suppress dendritic extension in Drosophila (13).

Although some studies suggest that small GTPases and GEFs may have different effects on axonal and dendritic growth (14–16), virtually no study has addressed the directional specificity of those signaling molecules for neurites with similar properties in the context of bidirectional growth. Here, we use the bipolar PLM neurons in *Caenorhabditis elegans* to investigate this question. The posteriorly located, bilaterally symmetric PLM neurons are two of the six mechanosensory touch receptor neurons (TRNs; 17). The PLM neurons have two sensory neurites that grow in different directions; both contain the MEC-4 transduction channel (18; Fig. S1) and have bundles of the 15-protofilament microtubules that are only present in the TRNs and are required for mechanosensation (19, 20). In addition, both neurites are capable of forming synapses if partners are physically close (21). In contrast to the PLM neurons, the four remaining TRNs (the two anterior ALM, the AVM, and the PVM neurons) are monopolar.

As part of a genetic screen for mutants with morphologically abnormal TRNs (to be reported elsewhere), we identified mutants defective in differential outgrowth of the two PLM neurites. In particular, we found that GEF proteins UNC-73/Trio and TIAM-1 promote neuronal extension towards the anterior and the posterior, respectively, and that the Rac subfamily GTPase CED-10/Rac1 is activated by TIAM-1, whereas CED-10 and MIG-2/RhoG act redundantly downstream of UNC-73. Moreover, the two pathways promoting growth in opposing directions antagonize each other to control neurite morphogenesis. Thus, our study suggests that intracellular signaling pathways confer directional specificity on neurite extension through the activation of distinct GEFs and Rac GTPases.

## Results

### UNC-73/Trio promotes anteriorly directed neurite extension

Previously we found that *unc-73/Trio* regulates axonal guidance in the TRNs (22). We isolated a new *unc-73* allele (*u908*; Fig. S2) in our recent screen and found that mutations in *unc-73* led to the specific shortening of TRN anteriorly-directed neurites (ANs). *unc-73* encodes multiple isoforms of a guanine nucleotide exchange factor that is homologous to mammalian Triple functional domain protein (TRIO). The different isoforms of UNC-73 contain a RacGEF domain, a RhoGEF domain, or both. One of the isoforms, UNC-73B, which contains only the RacGEF domain, specifically activates Rac pathways and affects axonal guidance, cell migration, muscle arm extension, and phagocytosis (9, 23–25). In contrast, a different isoform, UNC-73E, which contains only the RhoGEF domain, activates Rho GTPases and regulates both cell migration and locomotion (26, 27). We found that missense mutations *u908* and *rh40* and splice donor mutation *e936*, which all alter UNC-73B, resulted in shortened ANs from both the PLM and ALM. The PLM-AN often stopped before reaching the PVM cell body, the ALM-AN did not extend past the first pharyngeal bulb, and the AVM-AN stopped extending anteriorly after branching into the nerve ring (Fig. 1; the PVM-AN was not scored because it runs together with AVM-AN in the ventral cord and the AN ending could not be easily identified). Mutations *ce362* and *ev802*, which alter *unc-73E* did not cause similar defects (Fig. 1C).

**Figure 1.**
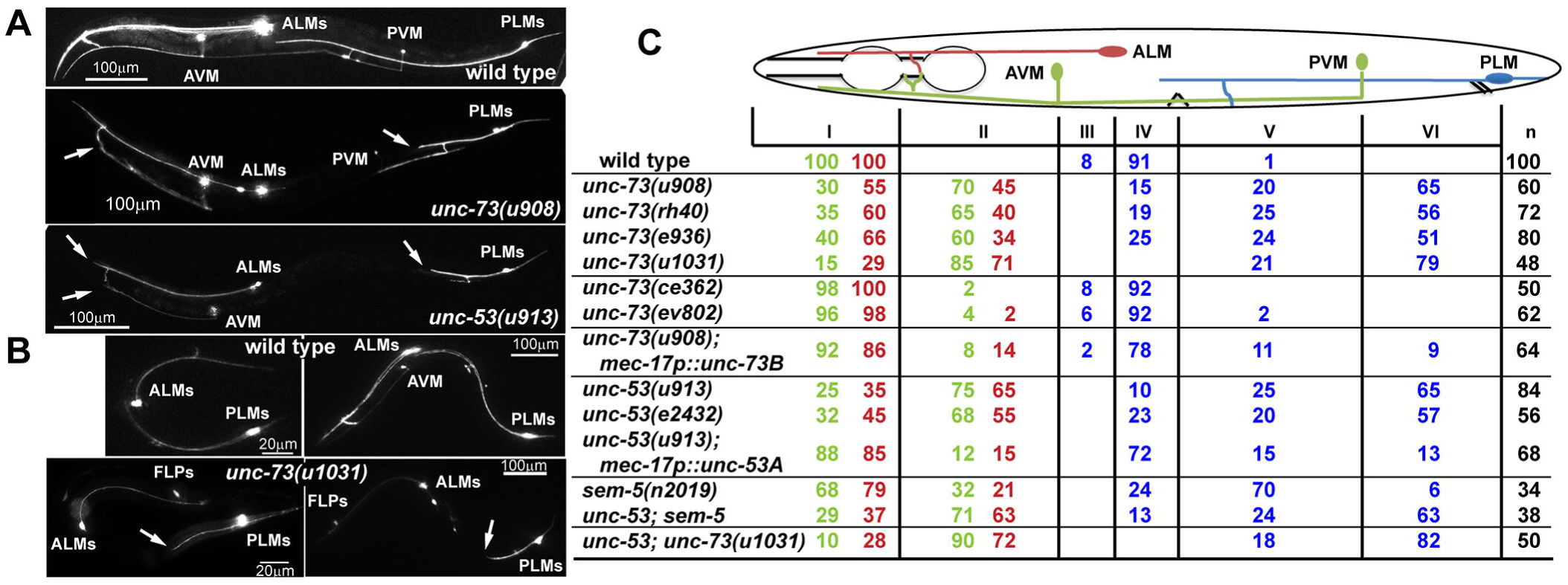
UNC-73 and UNC-53 promote anteriorly directed neurite extension. (A) Adult TRN morphologies, visualized by RFP expressed from the *mec-17* promoter, in wild-type, *unc-73(u908)*, and *unc-53(914)* animals. Arrows here and elsewhere in the figure point to the premature ends of the neurites. (B) TRN morphologies in wild-type or *unc-73(u1013)* animals at the second larval stage (left) and early adult stage (right). *mec-3p∷RFP* was used to label the TRNs and FLPs in *unc-73(u1013)* mutants. (C) The percentage of TRN anterior neurites terminating in specific zones (red numbers for ALM, green-yellow numbers for AVM, and blue numbers for PLM). The animal body is divided into six zones along the anterior-posterior axis. Zone I occupies the area from the nose to the posterior end of the first pharyngeal bulb; AVM and ALM cell bodies, vulva, PVM and PLM cell bodies define the boundaries from zone II to zone VI. Determined by a Chi-square test, results of *unc-53(e2432); unc-73(u1031)* were not significantly different (*p* > 0.05) from *unc-73(u1031)* single mutants; and the difference between *unc-53(e2432); sem-5(n2019)* and *unc-53(e2432)* was not significant.

Furthermore, we generated a *unc-73B* null allele, *u1031*, by creating a frameshift-causing 5-bp deletion in the second exon of *unc-73B* using CRISPR (clustered regularly interspaced short palindromic repeats)/Cas9-mediated genome editing (Fig. S2; 28). This null mutation produced sterile adults, which was not observed in animals carrying missense mutations, as well as a strong uncoordinated phenotype. The anteriorly directed neurite growth was also strongly suppressed in *unc-73B(u1031)* animals (Fig. 1C). Although the *u1031* mutation also affected *unc-73A*, which contains both RacGEF and RhoGEF domains, the morphological defects in the mutants could be rescued by specifically expressing the *unc-73B(+)* from the TRN-specific *mec-17* promoter (Fig. 1C). Therefore, we refer to *u1031* as an *unc-73B* allele in this context. The expression of *unc-73* in the TRNs was previously reported (29). Together, these results indicate that the Rac-specific GEF domain of UNC-73 promotes neurite extension towards the anterior in a cell autonomous manner.

van Haren *et al* (30) reported that TRIO complexed with NAV1 (neuronal Navigator 1) to regulate neurite extension by binding to the plus-end of growing microtubules in cultured mouse hippocampal neurons. UNC-53, the *C. elegans* homolog of Navigators, is also required for TRN outgrowth and guidance (31), and we found a specific shortening of TRN-ANs in *unc-53* mutants (Fig. 1C). Mutations in *sem-5/Grb2*, which encodes an adaptor protein for UNC-53/NAVs (32), also resulted in shorter TRN-ANs; the loss of *sem-5* did not enhance the phenotype of *unc-53*, suggesting they work together (Fig. 1C). The TRN morphological defects in the *unc-73 unc-53* double mutants were similar to *unc-73* null single mutants, suggesting that TRIO and NAVs act in the same pathway to promote neurite growth.

### TIAM-1 promotes posteriorly directed neurite extension

The growth of the PLM posterior neurite requires TIAM-1 (homolog of mammalian T-cell lymphoma Invasion And Metastasis 1), another Rac GEF (33). Both the *tiam-1* nonsense allele *u914* (R570*; isolated in our screen) and the deletion allele *ok772* caused significant shortening of the PLM-PN and overextension of the PLM-AN (Fig. 2A-C). Expression of *tiam-1(+)* cDNA from the TRN-specific *mec-17* promoter rescued these defects. Although the *tiam-1* promoter reporter was expressed in both ALM and PLM neurons (Fig. S3), we did not observe any morphological defects in *tiam-1*-deficient ALM neurons. However, overexpression of *tiam-1* in the TRNs caused the elongation of the PLM-PN and premature termination of the ALM-AN and PLM-AN in a dose-dependent manner (Fig. S4). These results suggest that TIAM-1 acts cell-autonomously to promote posteriorly-directed neurite growth and to suppress anteriorly-directed extension. Consistent with its positive effect on PLM-PN growth, TIAM-1∷GFP localized specifically to the distal region of PLM-PN and was not found in PLM-AN (Fig. 2E). Antibody staining against GFP in animals carrying the *mec-17p∷tiam-1∷GFP* transgene failed to detect any signal in PLM-AN either. Although we could not exclude the possibility that the amount of TIAM-1 in the anterior was below our detection limit, the effect of TIAM-1 loss on the overextension of PLM-AN is likely to be indirect. In contrast, UNC-73∷GFP, which rescued the *unc-73* mutant phenotype, was found as puncta along both PLM-AN and PLM-PN, suggesting that UNC-73 is ubiquitously present in the neuron but its growth-promoting activity is limited to the anterior neurites. Nevertheless, we cannot rule out that overexpression of UNC-73∷GFP caused it to be found in the PLM-PN.

**Figure 2.**
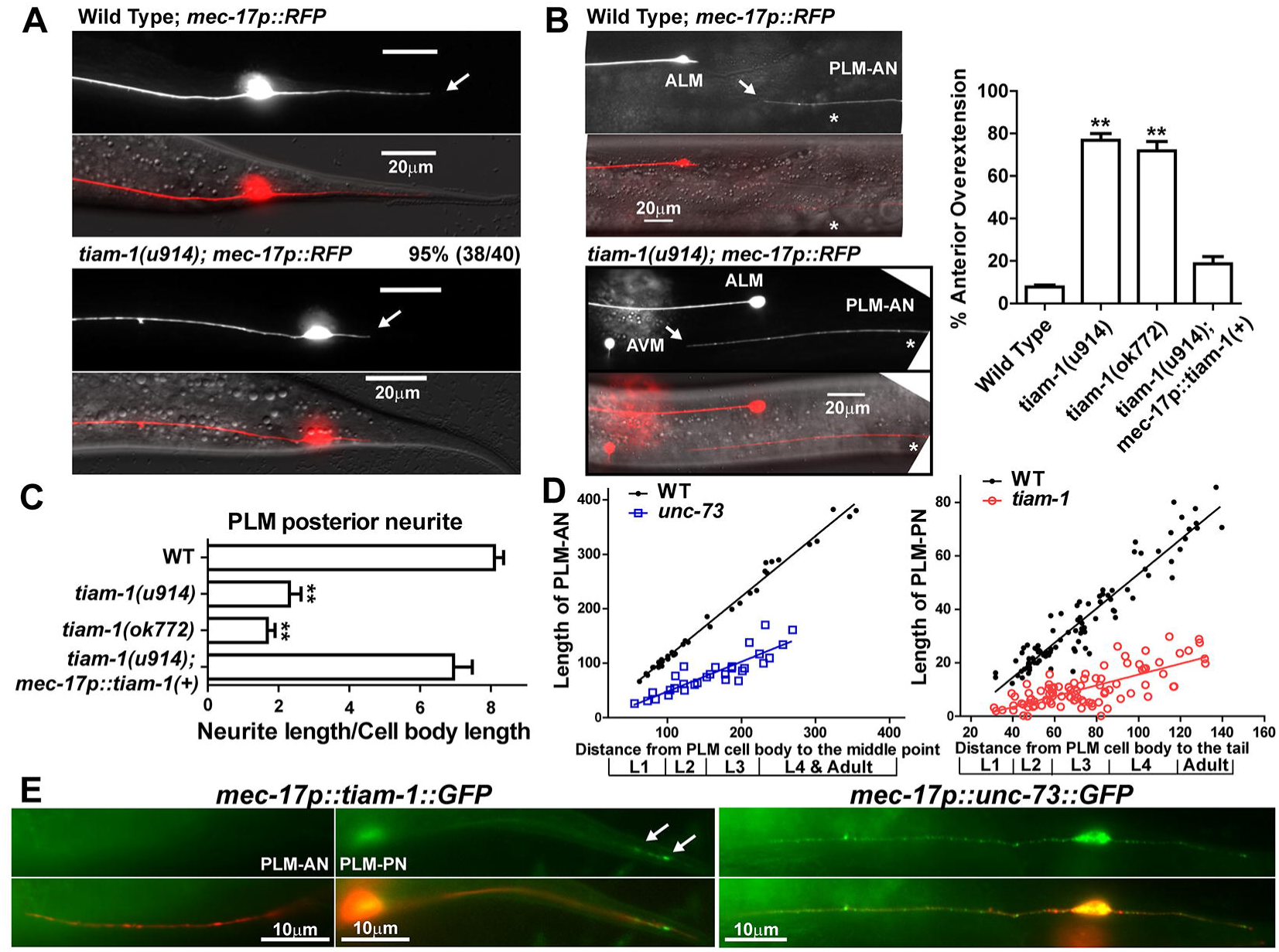
TIAM-1 promotes posteriorly directed neurite growth. (A) The shortening of PLM-PN in *tiam-1* mutants compared to wild-type animals. Arrows indicate the end of PLM-PN. (B) Overextension of PLM-AN in *tiam-1* animals, and the percentage of PLM-AN passing the ALM cell body. Asterisks in the images indicate the position of the vulva, and the arrows point to where PLM-AN terminates. (C) The length of PLM-PN in *tiam-1* animals. (D) The lengths (in μm) of PLM-AN and PLM-PN in wild-type, *unc-73(u1031)*, and *tiam-1(u914)* animals along different developmental stages. The distance from the PLM cell body to the middle point (the position of vulva in L4 and adults) or the tip of the tail was used as a measurement of developmental progress. Linear regression (solid lines) was applied to these data. The slope indicates the growth rate of neurites relative to the extension of the worm body. For WT PLM-AN, the slope = 1.1 ± 0.02 and r^2^ = 0.99 (goodness of fit); for *unc-73* PLM-AN, the slope = 0.55 ± 0.06 and r^2^ = 0.77; for WT PLM-PN, the slope = 0.65 ± 0.02 and r^2^ = 0.90; for *tiam-1* PLM-PN, the slope = 0.2 ± 0.02 and r^2^ = 0.55. (E) The localization of TIAM-1∷GFP and UNC-73∷GFP fusion proteins expressed specifically in the TRNs from L1 animals that carries the *mec-17p∷RFP* transgene. Arrows point to the position of TIAM-1∷GFP in the distal region of PLM-PN.

### UNC-73 and TIAM-1 regulate neurite elongation throughout development

The morphogenesis of PLM neurons can be divided into two phases: 1) the initial outgrowth of the two neurites and the establishment of a bipolar shape in the late embryonic stages; 2) the elongation of the two neurites throughout the larval stages. UNC-73 and TIAM-1 are required for the latter extension phase, because their mutants showed persistent neurite growth defects from the first larval stage to the adulthood (Fig. 2D); the two GEFs are unlikely to be involved in the initial neurite outgrowth, as the capacity of the neurons to form two opposing neurites was not affected in the mutants.

Heiman and Shaham proposed a “retrograde extension” mechanism for dendritic growth in *C. elegans* amphid neurons, showing that neurites can grow by anchoring their distal tips to specific locations and later being stretched as the cell body migrates away during larval development (34). UNC-73 and TIAM-1 are not likely to act similarly in PLM neurons, because the neurites in *unc-73* and *tiam-1* mutants showed substantial growth throughout development but with a much lower growth rate than the wild-type neurites (see SI results). Therefore, GEFs most probably regulate active neurite extension by affecting growth cone migration.

### CED-10/Rac1 promotes posterior neurite growth downstream of TIAM-1

Functionally, both UNC-73 and TIAM-1 activate Rac subfamily GTPases by stimulating the exchange of GDP for GTP; in *in vitro* nucleotide exchange assays, UNC-73B activates CED-10/Rac1 and MIG-2/RhoG (35), and TIAM-1 activates human Rac1 (33). *C. elegans* has three active Rac subfamily genes: *ced-10/Rac1*, *rac-2*, and *mig-2/RhoG*. Mutations in *ced-10*/Rac1 caused a significant reduction in the length of PLM-PN (Fig. 3A). This phenotype was enhanced when we expressed interfering hairpin RNAs targeting the third and fourth exons of *rac-2* from a TRN-specific promoter. [This RNAi experiment would also reduce the expression of a pseudogene *rac-3*, which is 98% identical to *rac-2* and appears to be a nonfunctional locus (36).] Therefore, TIAM-1, CED-10/Rac1, RAC-2 were needed for the production of the PLM posterior neurite. Moreover, *tiam-1; ced-10* double mutants have similar phenotype as *tiam-1* single mutants, supporting that Rac1 functions in the same pathway as TIAM-1 in promoting posteriorly directed extension (Fig. 3D). Mutations in another Rac gene *mig-2* or in the Cdc-42 genes (*cdc-42*, *chw-1*, and *crp-1*) did not alter PLM morphology (Fig. 3A).

**Figure 3.**
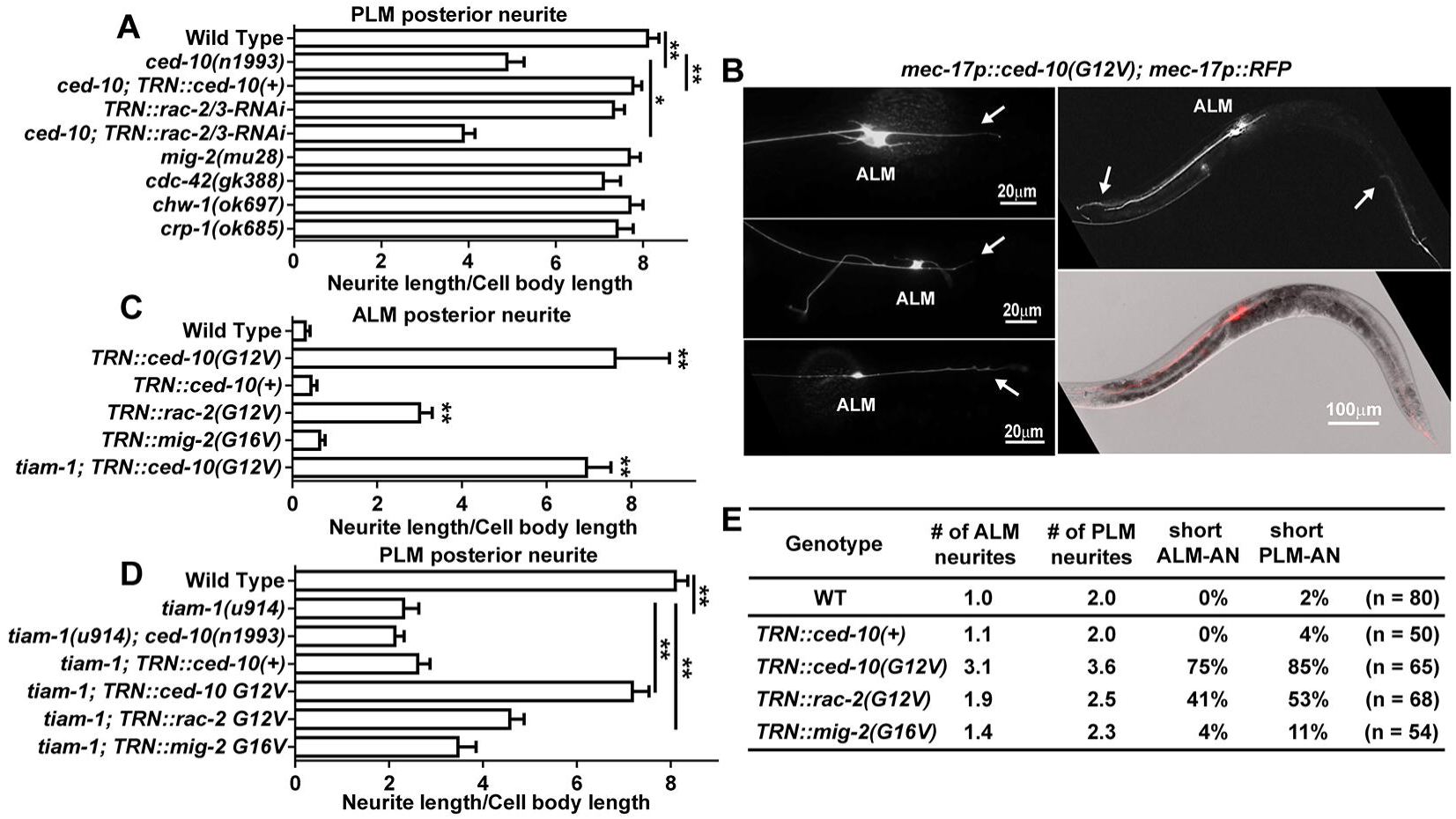
CED-10/Rac1 promotes posteriorly directed neurite outgrowth. (A) The length of PLM-PN in various strains. (B) TRN morphologies in animals expressing constitutively active CED-10(G12V) from the *mec-17* promoter. Left panels show morphological transformation of ALM neurons; arrows indicate the ectopic ALM-PN. Right panels show the premature termination of ALM-AN and PLM-AN; arrows point to the ends of the anterior neurites. (C-D) The length of ALM-PN and PLM-PN in various strains. (E) The average number of ALM and PLM neurites emanating from the cell bodies; the percentage of TRNs with shortened anterior neurites.

We next tested whether elevating Rac1 activity changed TRN morphology by expressing alleles of Rac genes [*ced-10*(G12V), *rac-2*(G12V), and *mig-2*(G16V)] encoding constitutively active proteins (36) in the TRNs. As in a previous study of Rac1 function in the PDE neurons (37), constitutively active CED-10 [CED-10(G12V)] induced the production of many lamellipodia-like protrusions from the plasma membrane, multiple short ectopic neurites from the cell body, and abnormal branches from the main TRN processes (Fig. 3B and E). Nevertheless, the most common (85%, n = 65) and dramatic phenotype resulting from the expression of CED-10(G12V) was the production of a long ectopic ALM posterior neurite (Fig. 3B and C). The ability of CED-10(G12V) to promote the generation of ALM-PN did not depend on the presence of *tiam-1* (Fig. 3C). In PLM neurons, hyperactive mutations in *ced-10* restored the PLM-PN in *tiam-1* mutants (Fig. 3D). These results suggest that CED-10 acts downstream of TIAM-1 to promote the growth of the posterior neurite. Constitutively active CED-10 also led to the shortening of both ALM-AN and PLM-AN (Fig. 3B, right panel), which indicates that the promotion of posteriorly directed neurite growth by TIAM-1 and Rac1 results in the inhibition of anterior extension.

Importantly, overexpression of wild type *ced-10(+)* did not cause a similar phenotype, indicating that the activity but not the level of CED-10 protein is crucial for posterior outgrowth. Constitutively active RAC-2 had a similar but weaker effect (Fig. 3), confirming that CED-10 is the major Rac GTPase regulating TRN neurite extension. Constitutively active MIG-2/RhoG, however, produced very little change in the TRN morphology in the wild-type animals and failed to rescue the outgrowth defects of *tiam-1* mutants (Fig. 3).

CED-10/Rac1-mediated regulation of neurite morphology appeared to be intrinsic, since the morphology of TRNs cultured *in vitro* was also controlled by Rac1 activity. Wild-type ALM and PLM neurons were morphologically distinct in culture: 78% of the ALM neurons had only one prominent neurite, whereas 76% of the PLM neurons had two neurites (Fig. S5A). When CED-10(G12V) was expressed in the TRNs, both ALM and PLM grew more than 2 neurites, although their average length was shorter than those of the wild-type cells (Fig. S5B-C). Thus, increased CED-10 activity resulted in the transformation of cell shapes and the ectopic growth of multiple neurites.

### CED-10/Rac1 and MIG-2/RhoG redundantly regulate anteriorly-directed elongation

*ced-10* and *mig-2* act redundantly downstream of *unc-73* in the regulation of cell division and migration (38). Although neither *mig-2* nor *ced-10* single mutants had anterior extension defects, we found that *ced-10; mig-2* double mutants had shorter TRN-ANs, most obviously in the PLM and AVM neurons (Fig. 4A). These morphological defects could be rescued by expressing either *ced-10(+) o*r *mig-2(+)*, suggesting that CED-10 and MIG-2 redundantly promote anteriorly directed extension.

**Figure 4.**
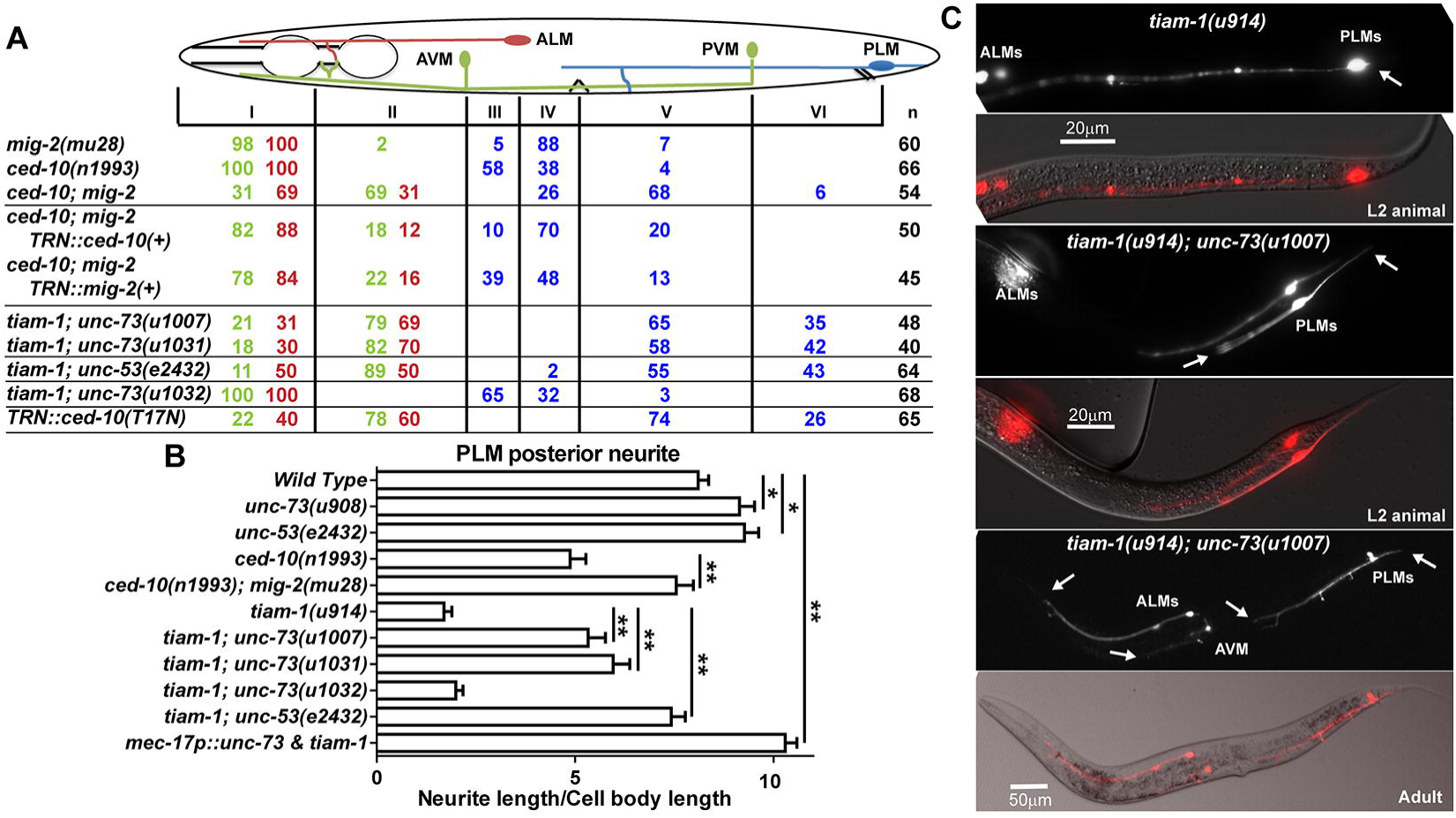
Mutations in *unc-73* suppress the shortening of PLM-PN in *tiam-1* mutants. (A) The percentage of TRN anterior neurites terminating in various zones (red numbers for ALM, green-yellow numbers for AVM, and blue numbers for PLM). Determined by Chi-square test, the results of *tiam-1(u914); unc-73(u1031)* animals were significantly different (*p* < 0.05) from that of *unc-73(u1031)* single mutants (data in Figure 1) for PLM anterior extension; difference between *tiam-1; unc-53(e2432)* and *unc-53(e2432)* was also significant. However, the results for ALM-AN and AVM-AN were not significantly different between *tiam-1; unc-73* double mutants and *unc-73* animals. (B) The length of PLM-PN in the strains indicated. (C) TRN morphology in L2 animals and adults of *tiam-1; unc-73* double mutants. Arrows point to neurite ends.

Unexpectedly, we found that mutations in *mig-2* suppressed the PLM posterior extension defects in *ced-10* mutants (Fig. 4B). PLM-PN in *ced-10; mig-2* double mutants was significantly longer than that in *ced-10* single mutants and was similar to the wild type. These results suggest that the suppression of anteriorly directed neurite extension in *ced-10; mig-2* double mutants simultaneously promoted outgrowth towards the posterior.

### Antagonism between anteriorly-and posteriorly-directed neurite extensions

Because mutations in *unc-73* and *tiam-1* differed in their effects on PLM neurite extension, we examined the PLM morphology in *tiam-1; unc-73* double mutants. Since *unc-73* and *tiam-1* genes are physically adjacent on the same chromosome, we recreated the *unc-73B* null mutations (the *u1007* and *u1031* alleles; Fig. S1B) in animals carrying *tiam-1(u914)* allele using the CRISPR method. The loss of *unc-73B* strongly rescued the posterior extension defects found in *tiam-1* single mutants, and the *tiam-1; unc-73* double mutants had less severe anterior growth defects compared to *unc-73* null animals, although PLM-AN was still significantly shortened compared to the wild type (Fig. 4; compared with Fig. 1). As controls, the *u1032* and *u1033* alleles, which affect *unc-73E*, had no effects on the Tiam-1 phenotype (Fig. 4A and B). Similarly, *tiam-1; unc-53* had restored PLM-PN but still shorter PLM-AN; again the PLM-AN defect was less severe in the double mutants than in *unc-53* single mutants (Fig. 4A and B). These results suggest that the loss of UNC-73 or UNC-53 causes downregulation of the growth signal for the anterior neurites and the derepression of posterior outgrowth, and the lack of TIAM-1 also partially reactivate the neurite growth towards the anterior. In the absence of both GEFs, PLM neurons adopt a default bipolar shape with two prominent neurites. Interestingly, mutation of *tiam-1* did not rescue the shortening of the anterior neurite in the monopolar ALM and AVM of *unc-73* animals, although *tiam-1* is expressed in these neurons (Figure 4A).

The mutual suppression between UNC-73 and TIAM-1 in PLM was not mediated by the changes in localization. TIAM-1∷GFP still localized to the distal region of the PLM-PN in *unc-73* mutants and the localization of UNC-73∷GFP in PLM-AN and PLM-PN were not affected in *tiam-1* mutants (Fig. S6). These data support the hypothesis that the posterior-specific TIAM-1 could functionally override the presence of UNC-73 in PLM-PN to promote posteriorly directed extension. In fact, when UNC-73 and TIAM-1 are both overexpressed in the TRNs, PLM-PN became elongated compared to the wild-type neurite despite the excessive amount of UNC-73 (Fig. 4B). Although most (92%; n = 85) of the TRN anterior neurites overexpressing the two GEFs had a normal length, moderate shortening of ALM-AN and PLM-AN were observed in the remaining 8% of the cells (Fig. S7).

The phenotype of *tiam-1; unc-73* double mutants was similar to that of *ced-10; mig-2* animals, suggesting that CED-10 acts downstream of TIAM-1 and that UNC-73 activates both MIG-2 and CED-10. The expression of a dominant-negative CED-10/Rac1(T17N) mutant (39) also reproduced the *tiam-1; unc-73* double phenotype presumably by sequestering both UNC-73 and TIAM-1 from their substrates (Fig. 4B). This result supports that CED-10 contributes to both anterior and posterior growth, although it is essential posteriorly and redundant anteriorly.

### UNC-73 and TIAM-1 also promote directional neurite elongation in other neurons

In addition to PLM neurons, UNC-73 and TIAM-1 also regulated the elongation of the anterior and posterior neurites, respectively, in ALNL/R and PLNL/R neurons, two pairs of bilaterally symmetrical neurons located at the tail region (SI results; Fig. S8). Antagonism between the two GEFs was also observed in these neurons. Moreover, in the bipolar BDU neurons, which are located in the anterior half of the animal, TIAM-1 specifically promoted the posteriorly directed neurite extension (Fig. S8). These results suggest that GEFs generally control the directional specificity of neurite extension along the A-P axis.

## Discussion

Ever since the observation that the activity of Rho family small GTPases is locally regulated in living cells (40, 41), researchers have searched for factors that control this localized activation. This issue is particularly important for understanding the development of complex neuronal structures, because a wide range of extracellular guidance molecules induce highly specific growth cone behaviors through only a handful of common Rho GTPases. How is this specificity achieved? Accumulated evidence, mostly from *in vitro* studies, suggests that neurons employ distinct GEFs to activate the same GTPases in response to different guidance cues, and the use of certain GEFs is often cell type-specific (4). Our *in vivo* study suggests that multiple GEF molecules can also function in the same neurons to confer directional specificity in the development of multiple neurites. Given that humans have 83 Rho family GEFs (42), we propose that specializing GEFs as signaling molecules that promote neurite growth in particular directions may be a general mechanism for the spatial control of Rho activation during neuronal development.

PLM morphology is regulated by Wnt signals, and different sensitivities to the same guidance cue may explain the use of distinct GEFs and Rac GTPases for the growth of the two opposing neurites. PLM-AN is repelled towards the anterior by Wnt proteins released at the posterior side of the cell body (21, 43). Although Rac1 is known to act downstream of Wnt signaling (44), our data suggest a novel link between Wnt signaling and the control of Trio and RhoG. PLM-PN grows against the gradient of the repulsive Wnt proteins, and this growth requires both the activation of intrinsic Hox proteins (45) and the attenuation of the extrinsic Wnt signals by Dishevelled (21). The specific involvement of these proteins on PLM-PN but not AN may lead to the selective employment of TIAM-1 for posterior extension.

A distinctive response to the same guidance cue in different cellular compartments has also been observed elsewhere (46, 47). Notably, semaphorin 3A repels the cortical axons of mouse pyramidal neurons but attracts their apical dendrites; this specific attraction is mediated by the asymmetric localization of soluble guanylate cyclase to the developing dendrites (34). Our findings suggest that this different responsiveness within the same cells can also be mediated by the local activation of distinct GEFs. In *C. elegans* PDE neurons, for example, TIAM-1 regulates growth cone extension *via* Rac GTPases responding to the activated netrin receptor UNC-40/DCC (33), whereas UNC-73 inhibits this extension (48). Importantly, lowering the levels of the *mig-2* and *ced-10* cause migrating cells to be repulsed rather than attracted to Semaphorin-1 and Plexin-1 (49), suggesting that the proper activity of Rho family GTPases is critical for the response to cues.

An unexpected finding of this research is that the two pathways promoting anteriorly and posteriorly directed growth antagonize each other. Mutations in *tiam-1*/GEF and *ced-10*/Rac1 caused significant shortening of the PLM-PN and overextension of the PLM-AN (Fig. 2 and 4), both of which were also observed in *egl-5* and *rfip-1* mutants (*egl-5* encodes a Abd-B-like Hox protein, and *rfip-1* encodes a recycling endosome-associated protein; 45). Therefore, suppression of the posterior extension appeared to allow overgrowth towards the anterior in general. On the other hand, *unc-73* and *unc-53* mutants had not only shorter PLM-ANs but also longer PLM-PNs (Fig. 4B), suggesting that suppression of anterior growth also led to overextension towards the posterior. The phenotype of the *tiam-1; unc-73* double mutants supports the mutual inhibition model, as the defects found in the single mutants were both alleviated in the double mutants. These data, however, also suggest that UNC-73 and TIAM-1 only regulate the directional bias of neurite extension and are dispensable for the general neurite growth capacity of the cells.

Based on the localization data (Fig. 2E), we hypothesize that the ubiquitously present UNC-73 biases neurite growth towards the anterior, promoting the growth of PLM-AN and suppressing the extension of PLM-PN. The PLM-PN-specific TIAM-1 counteracts UNC-73 activities, allowing the growth of a posterior neurite. However, the TIAM-1 localization does not explain the anterior overextension induced by the loss of *tiam-1* in both the wild-type and the *unc-73* animals. Indirect signaling may mediate such derepression. Alternatively, undetectably low amount of TIAM-1 in PLM-AN could limit anterior extension stimulated by UNC-73, whereas a high level of TIAM-1 in PLM-PN could override the effect of UNC-73 and drive posterior extension.

Although the mechanisms underlying the antagonism between the UNC-73 and TIAM-1 is still unclear, the two pathways may signal to block each other or simply compete for limited resources (50). In either case, uncontrolled growth of one neurite would restrain the growth of the other neurite. This reciprocal regulation is reminiscent of the competitive outgrowth of neurites during axonal specification (51) or the competition between the primary axons and their branches (52) and between axons and dendrites (53). Thus, our results support the notion that communication among the multiple neurites is necessary for a coordinated development of complex neuronal shapes.

## Materials and Methods

*C. elegans* wild-type (N2) and mutant strains were maintained at 20 °C, as previously described (54). Most strains were provided by the Caenorhabditis Genetics Center, which is funded by NIH Office of Research Infrastructure Programs (P40 OD010440). To create *unc-73B* and *unc-73E* null alleles, constructs that express CRISPR/Cas9 with sgRNAs targeting exon 2 and exon 21, respectively, were injected into either wild-type or *tiam-1(u914)* animals, using a previously described method (28). Most of the constructs were made using the Gateway cloning system (Life Technologies, Grand Island, NY); *ced-10*(*G12V*), *rac-2*(*G12V*), and *mig-2*(*G16V*) cDNA were kindly provided by Dr. Erik Lundquist. Transgenes *uIs31[mec-17p∷GFP]* III, *uIs115[mec-17p∷RFP]* IV, *uIs152[mec-3p∷RFP]* IV, and *uIs134[mec-17p∷RFP]* V were used to visualize TRN morphology, *uIs129[lad-2p∷GFP]* for ALN and PLN, and *uEx1005[ser-2p∷RFP]* for BDU neurons. To score the phenotype of premature termination of anteriorly directed TRN processes, the body of animal was divided into several regions and the percentages of TRN axons terminating at each specific region were calculated. Chi-squared test was used for the categorical data to find significant difference between different strains. The relative length of PLM neurites was calculated by dividing the length of the neurite by the diameter of PLM cell body along the A-P axis. For statistical analysis, ANOVA and a post hoc Tukey–Kramer method were used to identify significant difference between the sample means in multiple comparisons. Single and double asterisks indicated *P < 0.05* and *P < 0.01*, respectively. More details are provided in *SI Materials and Methods*.

## Acknowledgement

We thank Dan Dickinson, Erik Lundquist, and Yushu Chen for sharing reagents. This work was supported by NIH grant GM30997 to M.C.

